# Planting graphene quantum dots in PEGylated nanoparticles for enhanced and multimodal in vivo imaging of tumor

**DOI:** 10.1101/2022.03.27.486000

**Authors:** Hao Yan, Qian Wang, Jingyun Wang, Wenting Shang, Zhiyuan Xiong, Lingyun Zhao, Xiaodan Sun, Jie Tian, Si-Shen Feng, Feiyu Kang

## Abstract

Photoluminescent graphene quantum dots (GQDs) have recently attracted considerable attention for biomedical applications owing to the interesting physicochemical and photophysical properties, and prominent biocompatibility and biosafety. However, although much progress has been achieved in therapy and *in vitro* bioimaging, broad and crucial *in vivo* fluorescence bioimaging and synergistically working with other nanomedicine are very challenging and limited. Herein, we *in situ* implanted GQDs in the PEGylated layer of nanoparticles (NPs) via a bottom-up approach to obtain the NPs (core)-GQDs-PEG multifunctional nanoprobe (NPC-GQDs-PEG), which prolonged the blood circulation of GQDs more than four times and increased the tumor accumulation 7~8 times than GQDs used alone. Under assisted by the flexible microscope, the GQDs were successfully used for *in vivo* real-time monitoring of local NPs pharmacokinetics, *in vivo* multimodality imaging, and fluorescence imaging-guided tumor surgery. The approach of implanting GQDs in PEGylated nanomedicine and synergistic working provide a new strategy for *in vivo* biomedical applications of GQDs.

## Introduction

Graphene quantum dots (GQDs) are an emerging class of nanosized photoluminescent carbon nanomaterials with a characteristic size of sub-10 nm and outstanding physicochemical and photophysical properties (1–3). Compared with typical semiconductor quantum dots (semi-QDs), the GQDs show much better intrinsic biocompatibility and more easily renal clearance after the necessary thick surface coating for semi-QDs (4–6). In addition, GQDs have a smaller size (<2 nm of thickness) and molecular weight (dozens of kDa); therefore, they are more suitable for biological targeting and imaging than other quasi-spherical carbon dots (CDs) (usually hundreds of kDa) (4–5). Furthermore, the systemic cytotoxicity assessment has proved the excellent biocompatibility *in vitro* and *in vivo* and easy body clearance of GQDs (7–10). In the past decade, GQD has been explored extensively for biomedical application potentials, such as fluorescence bioimaging (5), magnetic resonance imaging (MRI) (11), drug delivery (12), and therapeutic applications (9–10, 13–14). From both fundamental and applied perspectives, the extremely stable fluorescence (FL) of GQDs is one of their most appealing characteristics (4). However, broad and crucial *in vivo* fluorescence bioimaging are still challenged and limited even some two-photon excited, and excellent near-infrared GQDs have been developed (15–17).

The low delivery efficiency of GQDs should be one of the main reasons for this after overcoming a series of physiological barriers *in vivo*. When the ultra-small size brings the rapid clearance via renal filtration, it concomitantly exhibits short blood-circulation half-lives and insufficient accumulation in targeted tissues (e.g., tumor) (18), which predominantly affect the effectiveness of GQDs *in vivo* (19). Therefore, surface engineering to lead a stealthy-to-sticky transition and functionalization with targeting or binding ligands is necessary for GQDs. However, these regular post-synthetic engineering strategies are strenuous and would affect the photophysical properties or even significantly quench the photoluminescence of GQDs (4–5, 20). Secondly, such restrictions of post-synthetic engineering would further hamper GQDs to composite with other nanomedicine to achieve synergistic effect and multimodality molecular imaging, which is especially needed in precision diagnostics (21–22). These limitations highlight the need to develop new strategies to formulate and composite GQDs to tune pharmacokinetics, enhance tumor accumulation and achieve specific bioimaging applications *in vivo*.

Among various approaches to developing ‘stealth’ long-circulating nanoparticles (NPs), the most widely used is poly (ethylene glycol) (PEG) grafting onto the surface of NPs (23), which dramatically decreases the undesirable clearance by the immune system *in vivo* (24). PEGylation of NPs can be easily functionalized with targeting ligands to further enhance their cell-binding ability (25). If the covalently attached PEG polymer is relatively less dense, macromolecules or proteins can penetrate the random coil PEG layer and reside in voids (26). Herein, we explored the strategy to implant GQDs *in situ* in PEGylated NPs via a bottom-up approach, generally formulate GQDs from small arene precursors (27), to form the NPs (core)-GQDs-PEG (targeted) multifunctional nanoprobe (NPC-GQDs-PEG) and then applied for *in vivo* bioimaging. In the PEG grafting layer voids, the GQDs was synthesized *in situ* from the precursor pyrene (C_16_H_10_) under mild conditions. Compared with GQDs alone, the seeding formulation, ‘stealth’ properties of PEG modification, and optimized design of NPC could extend the blood-circulation half-lives of GQDs by more than four times and increase the tumor accumulation by 7-8 times. Enrichment at the tumor of GQDs and following strong fluorescence signals was relied on to real-time monitor the local metabolism of NPs in tumors and major organs. With the versatility of NPs core, NPC-GQDs-PEG could achieve MRI and photoacoustic imaging (PAI) multimodality imaging *in vivo* and fluorescence imaging-guided tumor surgery. Seeding GQDs in PEGylated NPs makes *in vivo* bioimaging of GQDs more feasible and provides a new strategy to apply GQDs *in vivo*.

## Results

### Synthesis and characterization of NPC-GQDs-PEG nanocomposite

Graphene quantum dots are implanted into PEGylated nanoparticles to form NPC-GQDs-PEG through a hydrothermal condensation reaction, in which graphene-cell molecules pyrene (C_16_H_10_) as the precursor (Fig. 1a). First, the magneto-conjugated polypyrrole polymer (Fe_3_O_4_@PPy) nanoparticle core (NPC) was first synthesized following an in-situ oxidation polymerization method (28–29). The synthesized NPC is about 40~45 nm in diameter with a polymer shell of about 2~3 nm (Fig. 1b). Then, a long chain (10 KDa) amine-PEG-2-Deoxyglucose (PEG-2-DG) was added to react with the carboxyl groups of PPy to form PEGylation of NPC. The external 2-DG of PEG can significantly increase the NPs endocytosis of some cancer cells due to increased glucose consumption (29–30). Then, pyrene consisting of four peri-fused benzene rings was chosen as the precursor to growing GQDs in the PEG layer (Fig. 1a and Fig. S1), following a bottom-up procedure (31). The 1,3,6-trinitro pyrene, a derivative of pyrene through nitration treatment, penetrated PEG layer voids and in situ formed GQDs under hydrothermal treatment in the mixture of ammonia and hydrazine hydrate solutions (Fig. 1a).

**Figure 1.**
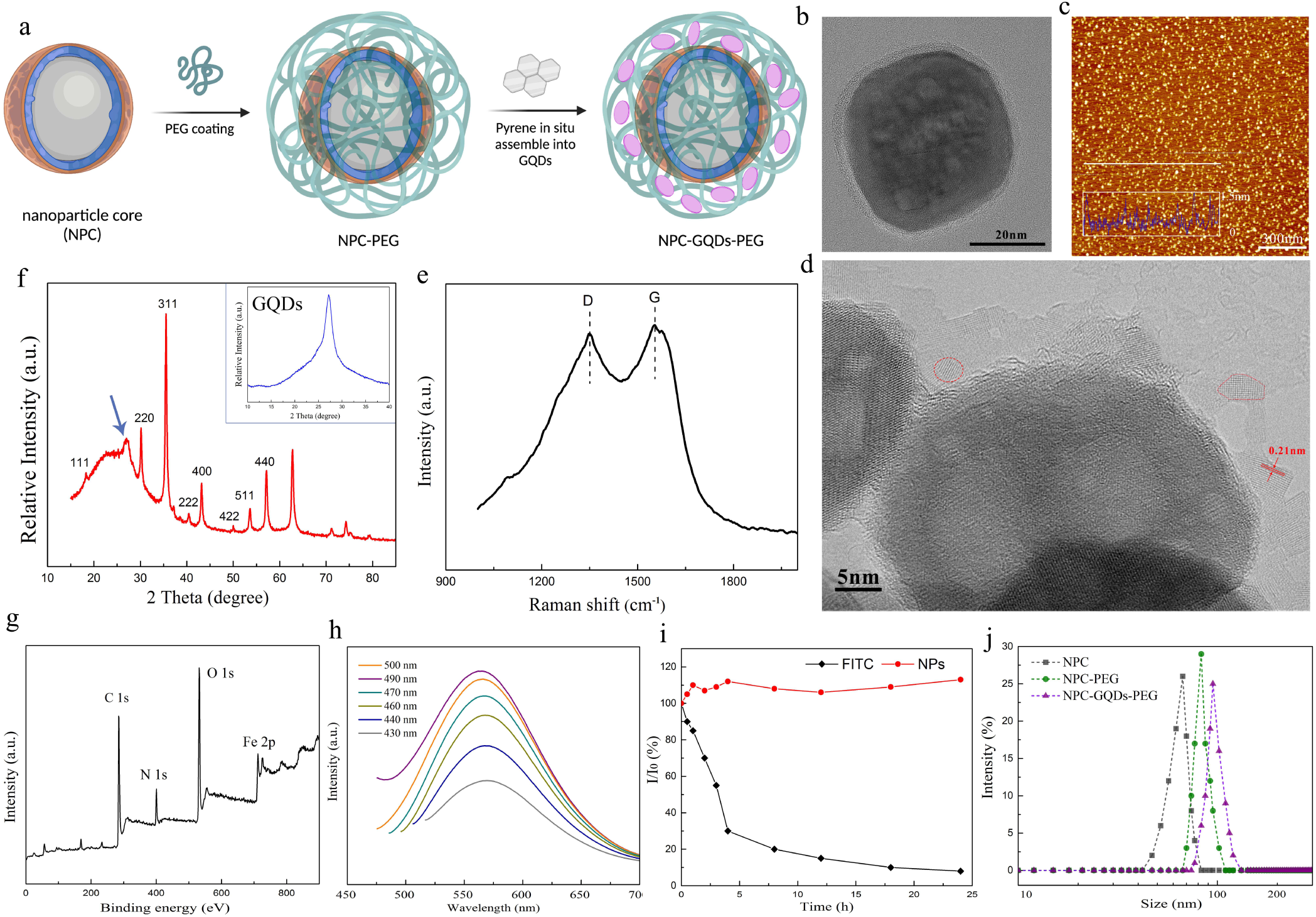
Preparation and characterization of NPC-GQDs-PEG nanoparticles. **a**, schematic of the process of GQDs *in situ* implanting into PEGylated NPs. **b**, HRTEM image of nanoparticle core (NPC). **c**, AFM image of pure GQDs (insets: height profile along the white line and height distribution). Scale bar, 300 nm. **d**, HRTEM image of NPC-GQDs-PEG nanoparticle, the red dotted lines mark single GQDs. **e**, Raman spectrum. **f**, XRD pattern of NPC-GQDs-PEG (insets: the XRD pattern of pure GQDs). **g**, Survey XPS spectrum. **h**, PL spectra of NPC-GQDs-PEG excited at different wavelengths (430 nm to 500 nm). **i**, Photostability of NPC-GQDs-PEG and organic dye FITC under 365-nm UV light (20 W). **j**, Hydrodynamic diameters of different nanoparticles measured by dynamic light scattering.

The pure GQDs, synthesized under the same condition, have a crystalline structure with the lateral size of 3~4 nm (Fig. S2) and thickness of 1.2 ± 0.48 nm corresponding to an average layer number of about 4 (Fig. 1c). The obtained NPC-GQDs-PEG nanocomposite has a uniform morphology and diameter of about 50 nm from the scanning electron microscope (SEM) image (Fig. S3). The high-resolution transmission electron microscopy (HRTEM) image depicts the three core-shell structures with crystalline Fe_3_O_4_ inner core, amorphous polymer shell, and outermost single-crystalline GQDs (Fig. 1d). The GQDs-PEG layer is about 5~10 nm consists of 1~3 single GQDs. And the GQDs have a single-crystalline structure with a spacing of 0.21 nm corresponding to that of graphene (100) planes. From the Raman spectroscopy (Fig. 1e), the intensity ratio of the ordered G band (1,590 cm^-1^) to the disordered D band (1,365 cm^-1^) is about 1.08, indicating the existence of carbon conjugate structure of NPC-GQDs-PEG. The X-ray powder diffraction (XRD) patterns (Fig. 1f) of NPC-GQDs-PEG retain the typical reflection peak of crystalline Fe_3_O_4_ cores, the broadened peak (2θ = 20°–25°) of polymer PPy and reflect *π* stacking of GODs with a peak near 2θ = 25°–27° which is close to that of pure graphite (3.34 Å). The survey X-ray photoemission spectroscopy (XPS) and the respective high-resolution spectrums (Fig. 1g and Fig. S4) not only reveal the presence of expected Fe-O (532.6 eV), C=C (284.5 eV), C-N (285.8 eV), C-O (286.9 eV) and O-H (531.6 eV) but also show the presence of NH_2_ (399.9 eV), indicating the GQDs in NPs are functionalized by NH_2_ which has been approved to locate at the edge sites of GQDs (31–32).

After GQDs embedding, the obtained NPC-GQDs-PEG shows excitation wavelength-dependent emission with a peak at around 560 nm under the excitation of 490 nm (Fig. 1h). The photoluminescence (PL) intensity of the NPC-GQDs-PEG is comparatively stable under 24 h continuous radiation with a 365-nm UV light (20 W), while the fluorescein isothiocyanate (FITC) dye rapid declines and finally attenuates to almost complete bleaching (Fig. 1i). Additionally, the NPC-GQDs-PEG nanocomposite inherits excellent near-infrared (NIR) absorbance properties, ranging from 800 to 1100 nm (Fig. S5a). Furthermore, after embedding GQDs, the NPC-GQDs-PEG also maintains the excellent super-paramagnetic properties and high saturation magnetization (Ms) of approximately 73 emu/g (Fig. S5b), which declines slightly from 84 emu/g of NPC (29). The inherited excellent optical-magnetic properties of NPC-GQDs-PEG could be used for magnetic resonance imaging (MRI) and photoacoustic imaging (PAI), as shown in Figures S5 c and d. During the process of PEG grafting and GQDs implanting, the zeta potential changed from −45 mV of NPC to −10 mV of NPC-PEG (2-DG), and finally increased to 10 mV of NPC-GQDs-PEG (amine-GQDs) or −15 mV (OH-GQDs). The pure GQDs were 30 mV of amine-GQDs and −25 mV of hydroxyl-GQDs, respectively (Fig. S6). Additionally, the hydrodynamic sizes of NPs increased from about 64 nm of NPC to 85 nm of NPC-PEG and finally rose to about 95 nm after GQDs embedding (Fig. 1j). The release curve of GQDs in NPC-GQDs-PEG over time shows the embedded GQDs can gradually release from the PEG layer, especially after shaking for days (Fig. S7). The strategy of implanting in the PEGylated NPs dramatically changed some physicochemical for GQDs.

### NPC-GQDs-PEG undergo slower blood clearance and higher tumor accumulation

Before being applied *in vitro* and *in vivo*, we first evaluated the intrinsic cytotoxicity of NPC-GQDs-PEG on MCF-7 human breast cancer cells and L929 murine fibroblast cells in culture using a Cell Counting Kit-8 (CCK-8) assay (Fig. S8). No significant cytotoxicity was observed from all two cell types for up to 72 h, even with high concentrations of 150 μg/ml, while exhibiting slight effect even at 300 μg/ml.

We then studied the blood clearance of intravenously injected NPC-GQDs-PEG and two surfaces functionalized GQDs (GQDs-OH and GQDs-NH_2_). The BALB/c mice bearing MCF-7 tumors in three groups (n=4) were intravenously injected with NPC-GQDs-PEG (20 mg kg^−1^), GQDs-OH (5 mg kg^−1^), and GQDs-NH_2_ (5 mg kg^−1^) respectively for the pharmacokinetics examinations. The overall clearance kinetics of GQDs-OH is like those of GQDs-NH_2_, but NPC-GQDs-PEG intuitively seems much slower in distribution and later clearance than pure GQDs (Fig. 2a, b, and c). Specifically, the distribution half-life time(t_α_,1⁄2) of two kinds of GQDs is about 0.4 h while about 1.46 h for NPC-GQDs-PEG and the elimination half-life time (t_β_,1⁄2) is about 4 h for GQDs while about 17 h (17.20 ± 5.16 h) for NPC-GQDs-PEG. Furthermore, the area under the concentration-time (AUC) curve is about 100~110 ID% h g^−1^ for GQDs, while the AUC of NPC-GQDs-PEG is as high as 225 ID% h g^−1^ in the first 12 h (AUC_0-12h_). The difference for this AUC widens as time goes on. The difference between NPC-GQDs-PEG and GQDs is about 200 ID% h g^−1^ at 24 h and reaches about 270 ID% h g^−1^ at 36 h while about 120 at 12 h (Fig. 2d). After embedding in the PEGylated NPs, GQDs are more persistent in the blood circulation than GQDs alone *in vivo*.

**Figure 2.**
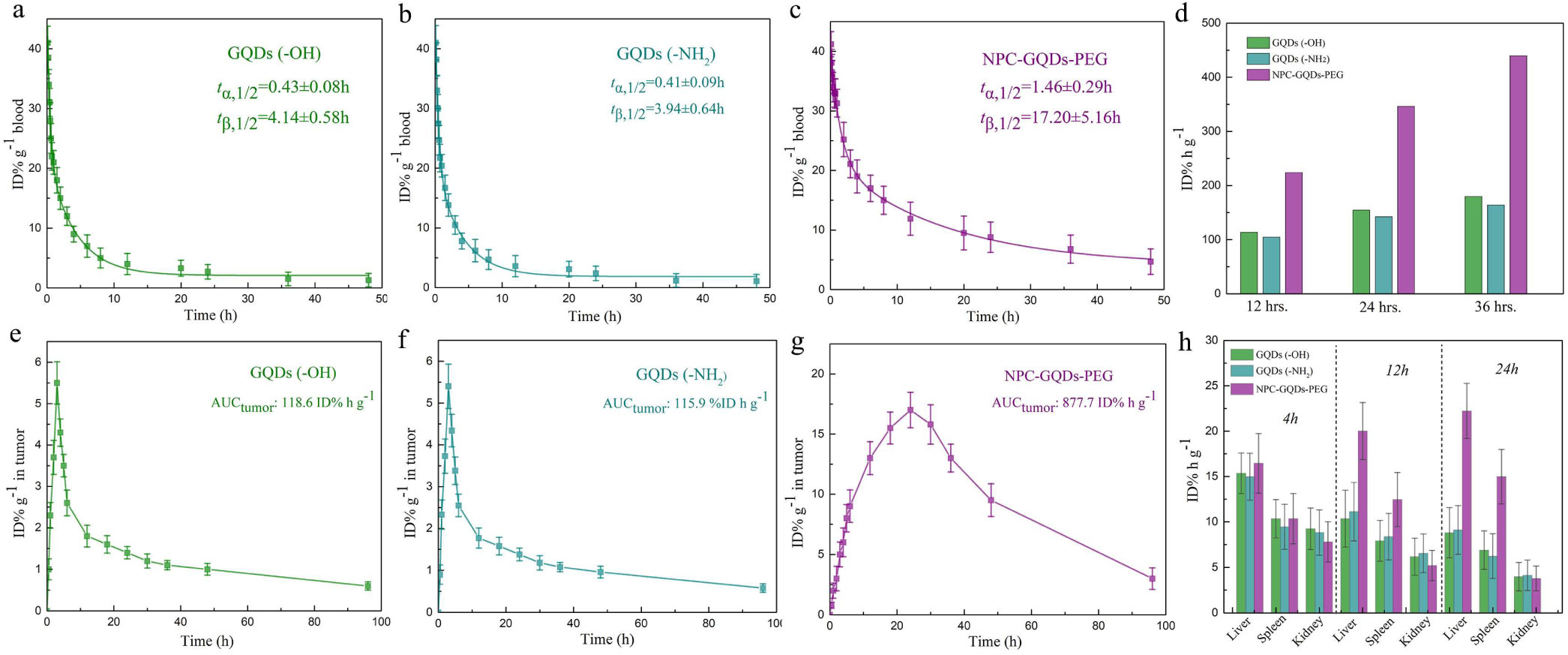
In vivo blood clearance and biodistribution of NPC-GQDs-PEG. **a-c**, Blood clearance kinetics in terms of percent injected dose (ID%) per gram blood for the GQDs (a-b) and NPC-GQDs-PEG (c) in mice after tail-vein injection of different NPs. GQDs-equivalent dose: 100 μg, and NPC-GQDs-PEG dose: 500 μg. **d**, Quantitative area under the concentration-time (AUC) pharmacokinetics curve of GQDs and NPC-GQDs-PEG at 12 h, 24 h, and 36 h after intravenous injection. **e-g**, Biodistribution profiles of GQDs (e-f) and NPC-GQDs-PEG (g) in xenograft MCF-7 tumors, in terms of percent, injected dose (ID%) per gram tumor after intravenous injection. The area-under-the-curve of tumor (AUCtumor) was calculated by accumulating the areas of the small trapezoids (between two-time points) under the curve. **h**, Quantitative biodistribution analysis of the GQDs and NPC-GQDs-PEG in the major organs at different time points post-injection. Data are mean ± s.d. of biological replicates (n = 4 mice per group).

It’s widely documented that more extended blood circulation could provide more chance for NPs to reach the tumor through the enhanced permeability and retention (EPR) effect (33). The concentration profiles of NPC-GQDs-PEG and GQDs in tumors to times were further investigated (Fig. 2e, f and g). The area-under-the-curve of tumor (AUC_tumor_) is 877.7 ID% h g^−1^ for NPC-GQDs-PEG while the AUC_tumor_ is just 118.6 ID% h g^−1^ and 115.9 ID% h g^−1^ for GQDs (-OH) and GQDs (-NH_2_) respectively. Following the previously reported method (18, 34), we further calculated the tumor delivery efficiency of NPs (Fig. S9, Supplementary equation 1-1 and 1-2) by dividing the AUC_tumor_ by the time of the endpoint (t_end_) and then multiplying the average tumor mass (m_tumor_). The tumor delivery efficiency of NPC-GQDs-PEG is 3.12%; contrastively, GQDs(-OH) and GQDs(-NH_2_) are just 0.42% and 0.41%, respectively. The strategy of embedding GQDs in PEGylated NPs could increase the tumor delivery efficiency of GQDs about 7.5 times in the xenograft breast cancer model. Further quantitative biodistribution analysis of NPC-GQDs-PEG and GQDs in the mice was conducted by measuring the samples of major organs at different time intervals post-injection. Compared with GQDs alone, NPC-GQDs-PEG is gradually enriched in the liver and spleen due to reticuloendothelial system absorption and long blood-circulation half-lives. The pure GQDs should be rapidly and efficiently excreted through the urinary and eliminated from the body owing to the ultrasmall size (35).

### *In vivo* real-time tracking NPC-GQDs-PEG in local tumors and organs

Real-time tracking the dynamics of systemically administered drugs in tumor microenvironments *in vivo* is believed vital importance for evaluating the efficacy of medicines (36–37). We further explored the real-time and long-term tracking of NPC-GQDs-PEG in the tumor microenvironment based on the excellent photostability of GQDs. Fibered confocal fluorescence microscopy contains two different kinds of laser probes to visualize NPC-GQDs-PEG on a microscopic level *in vivo* (Fig. 3a and Fig. S10). The local pharmacokinetics of NPs were monitored and assessed in tumors and some major organs (live, spleen, and kidney) in Balb/c mice bearing xenograft MCF-7 tumors. We first evaluated the intracellular uptake and distribution of NPC-GQDs-PEG *in vitro* and found that the NPs were efficiently uptake by MCF-7 cells and distributed around the nucleus in the cytoplasm (Fig. 3b). The GQDs implanting seems no noticeable effect on the targeting effect of 2-DG PEG to cancer cells.

**Figure 3.**
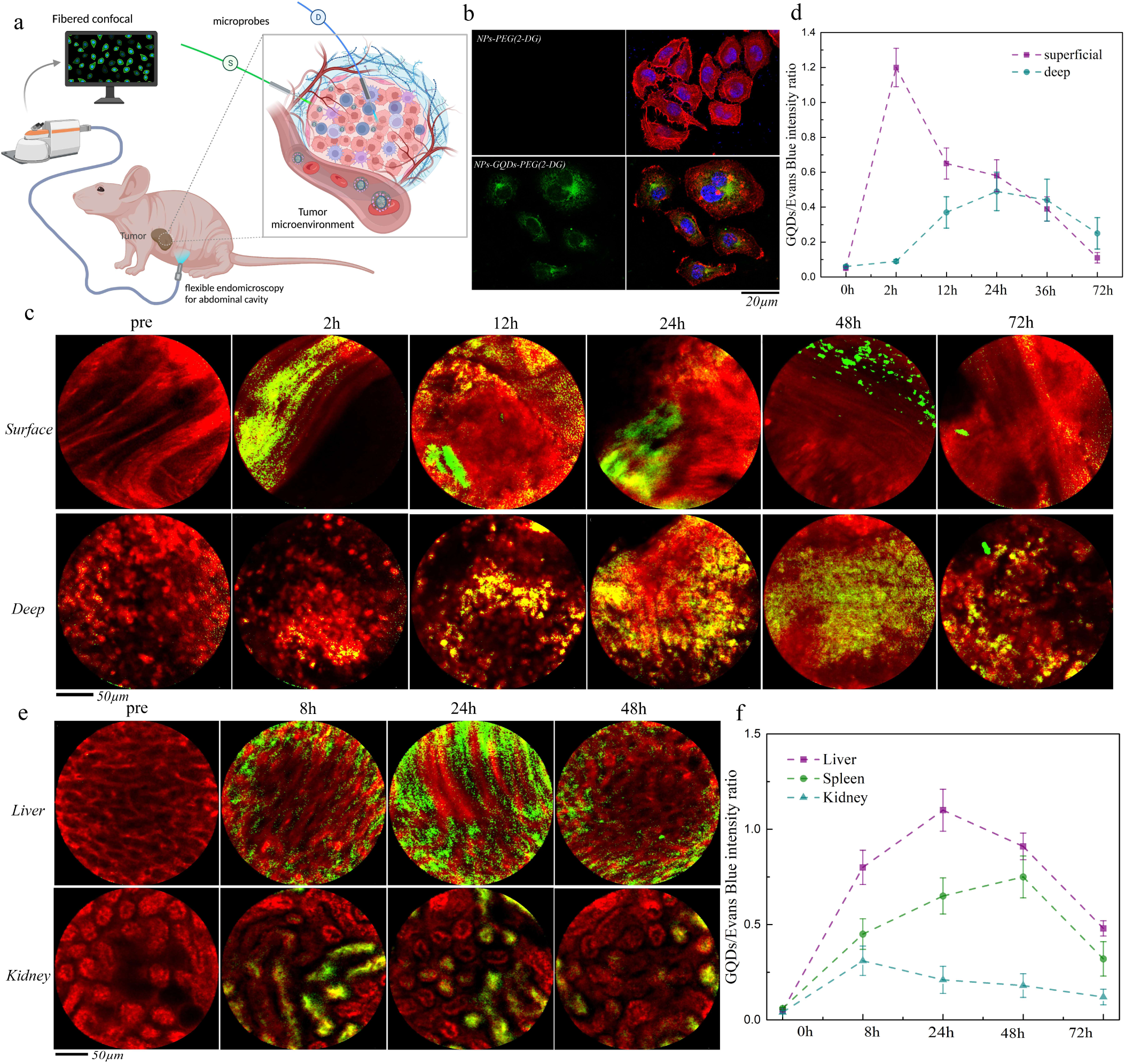
Real-time monitoring of the NPC-GQDs-PEG in tumor site and major organs. **a**, Schematic illustration of the fibered confocal fluorescence microscopy (FCFM) imaging for tumor tissues and major organs in the abdominal cavity. For tumor tissue, the optical fiber probes were inserted into the superficial (S-probe) and deep tumor tissue (D-probe), respectively. A small incision was made in the tumor epidermis, and the S-fiber (about 0.3 mm of tip diameter) was attached to the surface of the tumor tissue. The D-fiber (about 2.6 mm of tip diameter) penetrated the deep tumor after carefully removing a portion of the surrounding tissues. The D-fiber penetrated the abdominal cavity for major organs through a pre-operated small incision in the belly. **b**, Confocal fluorescence microscopy images of MCF-7 cells treated for four h with NPC-GQDs-PEG, cytoskeleton was stained with Rhodamine B (red)-labeled phalloidin, the nucleus was counterstained with (blue) 4’, 6-diamidino-2-phenylindole (DAPI), and the NPC-GQDs-PEG were green. **c**, *In vivo* monitoring of the accumulation, diffusion, and biodistribution of NPC-GQDs-PEG in tumor tissues using FCFM. NPC-GQDs-PEG showed green fluorescence (excited at 480 nm) while Evans blue (red, excited at 660 nm) was used as the vascular contrast agent. **d**, Quantification of the fluorescence intensity of NPC-GQDs-PEG in superficial and deep tumor tissue at different time points. Values are expressed as the ratio of fluorescence intensity from GQDs and Evan’s blue as mean ± SD (n=4). **e**, *In vivo* monitoring of the biodistribution and metabolism of NPC-GQDs-PEG in live and kidney using FCFM, Evans’s blue (red, excited at 660 nm) works as the vascular contrast agent. **f**, Quantification of the fluorescence intensity of NPC-GQDs-PEG in living, spleen, and kidney at different time points. Values are expressed as the ratio of fluorescence intensity from GQDs and Evans blue as mean ± SD (n=4).

For tumor imaging, one optical fiber with about 0.3 mm of tip diameter attached to the tumor surface through a small incision in the tumor epidermis, and another optical fiber further penetrated the deep tumor tissue (Fig. 3a). Evans blue dyes were used as a vascular contrast agent and carefully pre-injected via tail vein 4 hrs. Before imaging, therefore, in addition to staining blood vessels, it would also leak to surrounding tissues to some degrees. Then, the NPC-GQDs-PEG solution (25 mg kg^−1^) was intravenously injected into the mice. As shown in Figure 1c, the superficial blood vessels are visible, and no clear GQDs signals in the superficial or deep tissue (threshold was set relatively high to exclude the background interference) before injection. Two hours post-injection, the strong GQDs signals were found in the vasculature of superficial tissue, while the signals in the deep tissue were still very weak. From 2 h to 24 h post-injection, NPC-GQDs-PEG gradually enriched in the tumor vessels, and more and more NPs started to infiltrate the surrounding tissue and penetrate the deep tissue. As longer post-injection, NPC-GQDs-PEG were largely metabolized from the tumor, especially for the superficial vasculature and tissues from 48 h to 72 h. In contrast, the clearance of NPs in deep tissue was much slower, and many residues remained even 72 h post-injection, which is mainly due to the incompleteness of vasculature and higher tissue pressure. The quantitative fluorescence calculation further confirms these results (Fig. 3d) and exhibits that NPC-GQDs-PEG ‘leave’ the tumor tissue more slowly than ‘enter’ the tumor, incredibly slowly for the deep tumor tissue.

The capability of real-time and long-term tracking NPC-GQDs-PEG *in vivo* can be expanded to other organs. To allow the optical fiber to penetrate deep into the abdominal cavity, we made a small opening (about 4 mm wide) in the belly. We then inserted the optical fiber to probe the liver, spleen, and kidney (Fig. 3a). After intravenous injection, NPC-GQDs-PEG were gradually accumulated in the liver and spleen (Fig. 3e, Fig. S11, and Fig. S12) and reached the highest concentration at 24 h and 48 h for live and spleen, respectively (Fig. 3f). And then, the signals of GQDs were gradually decreased, indicating that the NPC-GQDs-PEG were slowly metabolized from the liver and spleen. This is consistent with the typical characteristics of distribution and metabolism of many similar-sized NPs *in vivo* (38). Interestingly, the signal of GQDs in the kidney had not disappeared with the almost complete clearance of NPC-GQDs-PEG from the blood 48 h and 72 h post-injection (Fig. 3e, 3f and Fig. S11). These residual GQDs signals may come from the gradual release of GQDs from NPC-GQDs-PEG over time *in vivo*. Therefore, the embedded GQDs have enabled real-time tracking of the dynamics of systemically administered NPs in tumor microenvironments and major organs.

### Multimodality imaging and fluorescence imaging-guided tumor surgery

The exponentially increasing accumulation of GQDs in tumors encourages further applications *in vivo*. Fluorescence molecular imaging (FMI) has been developed as a surgery guidance tool to precisely distinguish tumor margins from normal tissues intra-operatively (39–40). We further explored the NPC-based MRI-PAI multimodality imaging for guidance of preoperative surgical plan and then GQDs-based FMI-guided breast cancer surgery. The biodistribution of NPC-GQDs-PEG in the whole body and local tumor was directly observed by the MRI (Fig. 4a and Fig. S13). Considerable signals were found 12 h after intravenous injection and reached the maximum 24 h post-injection and then gradually decreased until 96 h post-injection. The 3D and 2D PA imaging (Fig. 4b and Fig. S14) showed the massive accumulation and wide distribution of NPs in the whole tumor 24 h post-injection. Additionally, 24 h after injection of NPC-GQDs-PEG, the obtained MR images provide precise localization of the tumor in the entire body with excellent soft-tissue contrast. The PA images of the whole tumor provide more information inside the tumor with rich optical contrast and high spatial resolution.

**Figure 4.**
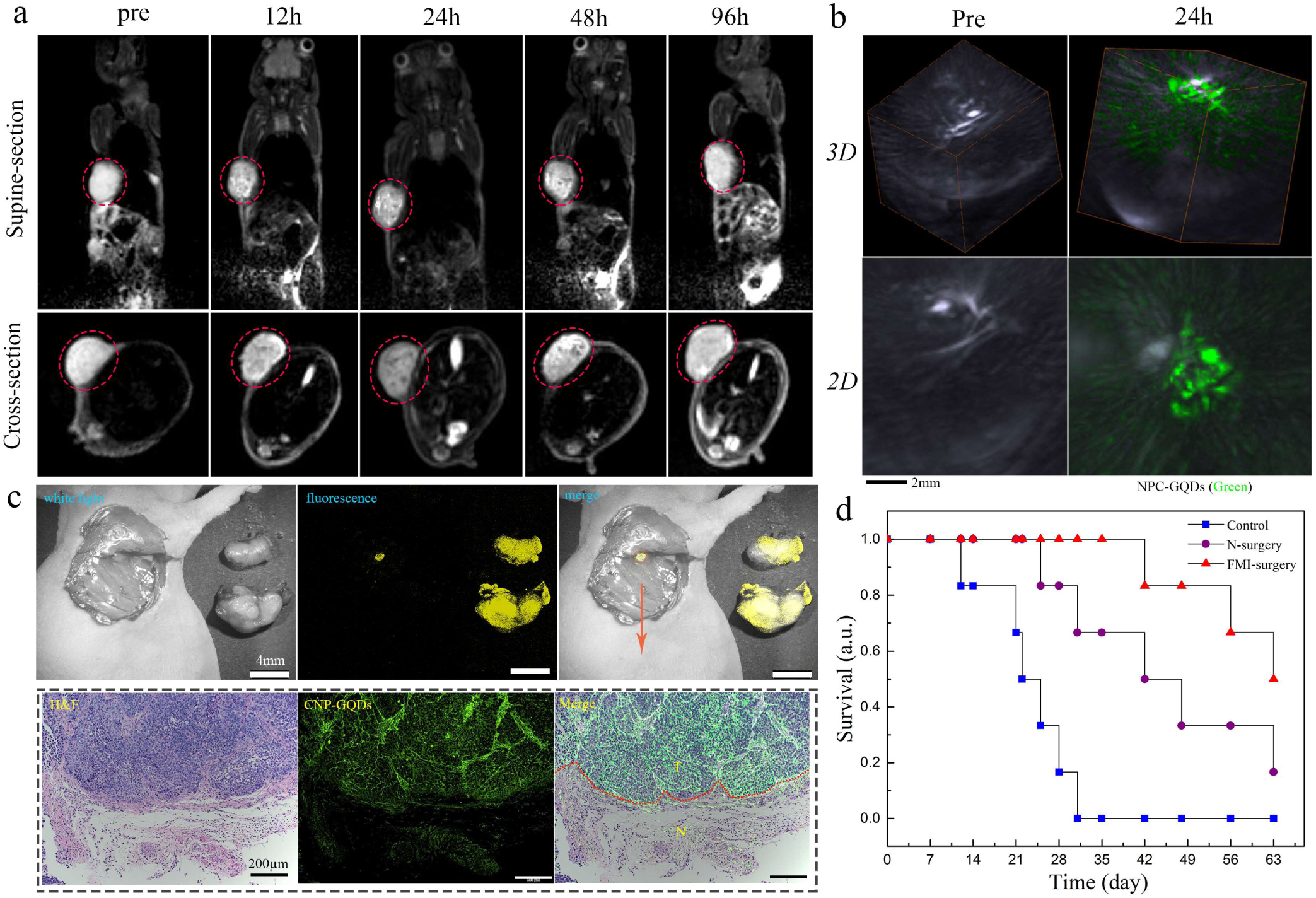
Multimodality imaging and FMI-guided tumor surgery using NPC-GQDs-PEG. **a** Representative *T*2-MR images of MCF-7 tumor-bearing mice at different time post *i*.*v*. Injection, including supine section and cross-section visions. **b**, 3-dimensional (3D) and 2D PA images of the tumor before and 24 h after *i*.*v*. Injection. The bright area is the tumor blood vessels which are rich in hemoglobin. The green signals correspond to the distribution of NPC-GQDs-PEG. **c**, Fluorescence imaging-guided intraoperative location, and removal of a tumor from mice bearing MCF-7 tumor (upper). Under the guidance of FMI, this residual microtumor was removed until no fluorescence signal was found. The subsequent histological analysis of the resected tumor foci (down). The tissue section was stained with hematoxylin and eosin stains (H&E), and the NPs were green due to GQDs. The boundary between the tumor and the normal tissue is visible (indicated with a red dashed line). **d**, Survival curves of the breast cancer-bearing mice after different kinds of treatment: no treatment (control), normal surgery (N-surgery), and fluorescence imaging-guided surgery (FMI-surgery). Error bars are the standard deviation of 6 mice per group.

Based on the comprehensive diagnosis information provided by the dual-modality imaging, we further explored the FMI-guided surgery to define the margins of tumor and detect residual microtumors. 24 h post-injection of NPC-GQDs-PEG, two experienced surgeons were assigned to try their best to completely remove the tumor tissues while minimizing the surgical trauma (Fig. S15). High-resolution intraoperative fluorescence images correlated with digital photographs were observed (Fig. S16). After surgeons under visual inspection completed the tumor resection, a small residual focus was still found on the resection bed (Fig. 4c, upper), which was so small that surgeons could not distinguish it (~0.5 mm). Then, the small focus was removed until no fluorescence could be detected in the resection bed. The last resected tissues contain prominent tumors tissues, and the tumor margins can be clearly distinguished from fluorescence image (Fig. 4c, down). After FMI-guided surgery, the survival rates of mice were continuously assessed and compared between different groups (n=6 in each group). The survival rates of mice in the FMI-guided group (FMI-surgery) are significantly higher than the normal surgery (N-surgery) and control groups (Fig. 4d). The 50% survival rate is 63 days for FMI-surgery, 42 days for N-surgery, and 23 days for the non-treatment group. Therefore, the FMI guidance of GQDs can tremendously improve the effectiveness of surgery.

## Discussion

The scheme of *in situ* implanting GQDs in the outer layer of PEGylated nanoparticles following the bottom-up process provides a new strategy for bioimaging applications of GQDs. PEGylation of NPs is the most widely used approach to mask their surface to obtain ‘stealth NPs’ *in vivo* by reducing blood protein adsorption and decreasing the clearance by the mononuclear phagocytic system (MPS) (24). Therefore, embedding GQDs into the voids of the PEG layer of NPs is a generic method. This approach also allows GQDs to inherit the rational design and physicochemical properties of PEGylated NPs, such as size, shape, charge, and targeting strategy, which will dominate the fate of GQDs *in vivo*. The final hydrodynamic size of about 95 nm, the near-neutral charge, and the targeting terminal 2-DG of NPC-GQDs-PEG significantly increase the circulation lifetime of GQDs and enable more GQDs to overcome level-by-level biological barriers to reach tumors. Furthermore, enough accumulation of GQDs in the tumor site allows further applications *in vivo*.

The embedding of GQDs into PEGylated NPs can further provide a multifunctional platform for multimodality molecular imaging, which can overcome the intrinsic limitations of each imaging method and provide comprehensive and precise diagnostic information. GQDs-based fluorescence molecular imaging can reveal the real-time pharmacokinetic and dynamic biodistribution of NPs in local with high-sensitivity. At the same time, NPs-based MRI provides the full-body imaging with high soft-tissue contrast and PAI reveals tumor pathophysiological status with high spatial resolution and deep penetration. Interestingly, the embedded GQDs could gradually release from PEGylated NPs and finally be cleared from the body, which maintains the excellent long-term biosafety of GQDs. Of course, the penetration depth of bottom-up grown GQDs here is very limited owing to the visible emission PL. However, there is hope to achieve near-infrared GQDs-embedded NPs-PEG with the development of bottom-up syntheses. Therefore, such a strategy of promoting strengths and avoiding the weakness of GQDs will broaden the *in vivo* bioimaging horizons of GQDs.

## Materials and Methods

### Synthesis and modification of nanoparticle core (NPC)

NPC is the core-shell Fe3O4@polypyrrole (PPy) optical-magneto nanoparticles. Fe3O4 nanoparticles of about 40 nm were first synthesized by a modified solvothermal method following our previous procedures in an ethylene glycol/diethylene glycol (EG/DEG = 1:5) binary system (41). Then, the conjugated polymer PPy was coated on the surface of the magnetic core following an *in-situ* polymerization method (28). Pyrrole-COOH was chosen as a monomer that can be easily grafted with amino-terminal poly (ethylene glycol) (PEG). The obtained NPC was then grafted with amine-PEG-2-Deoxyglucose (PEG-2-DG). Specifically, 5 mg long-chain NH2-PEGn-GLUC (MW = 10000) dissolved in 1 ml of dimethyl sulfoxide (DMSO) and then mixed with 5 ml NPC solution (500 μg/ml). After vertexing for 30 min, 5 mg N- (3-dimethyl aminopropyl)-N-ethyl-carbodiimide hydrochloride (EDC) (dissolved in DMSO) and 2.5 mg N-hydroxysulfosuccinimide (Sulfo-NHS) were added and reacted for 12 h in the dark with continuous stirring. The final particles were centrifuged (15,000 × *g* for 30 min) and washed three times for further use.

### Synthesis of NPC-GQDs-PEG

1 g pyrene was added to 80 ml HNO3 under refluxing and stirring for 12 h to obtain trinitropyrene and then diluted with deionized water filtered through a 0.22 μm microporous membrane to remove the acid. Meanwhile, 2 ml NPC-PEG solution (1 mg/ml) was dispersed in 30 mL ammonia (0.4 M)/hydrazine hydrate (1.5 M) mixed solvent. The above 1,3,6-trinitropyrene (0.25 g) was then added into 30 ml prepared NPC-PEG (50 μg/ml) ammonia (0.4 M)/hydrazine hydrate (1.5 M) solution. After further sonicated for 30 min, the suspension was transferred to a poly(tetrafluoroethylene) (Teflon)-lined autoclave (40 ml) and heated at 100 °C for 5 h. The resulting product was collected with the help of a magnet and purified with repeated centrifugation (20 000 g for 30 min) five times. The purified NPC-GQDs-PEG were dried at 80 °C for further characterization and property measurement. The pure GQDs-NH2 and GQDs-OH were synthesized following the above procedure except using different reaction mediums: 0.2 M NaOH solution for GQDs-OH while 0.4 M ammonia and 1.5 M hydrazine hydrate for GQDs-NH2.

### Characterization of GQDs and NPC-GQDs-PEG

The size and morphology of NPC-GQDs-PEG were characterized by SEM (JSM-7001F) and TEM (Hitachi HT7700). High-resolution TEM (HRTEM) images were obtained using a JEM 2100F transmission electron microscope. AFM images of GQDs were taken using an SPM-9600 AFM. Raman spectra were recorded on a Horiba (LabRAM HR Evolution) in plus laser Raman spectrometer with λexc= 785 nm. XPS spectra were performed on Escalab 250Xi XPS spectrometer (Thermo Fisher). Powder X-ray diffraction (XRD) patterns were measured on a Bruker D8 Advance X-ray diffractometer using a Cu-Kα X-ray source. A dynamic light scattering instrument (Malvern Zetasizer Nano-ZS, 633 nm) measured the hydrodynamic size and Zeta potential. The Fluorescence emission spectrum was measured by fluorescence Spectro-fluorometry (F-7000, Hitachi) at room temperature. The release curve of GQDs in NPC-GQDs-PEG over time was calculated by measuring the residual fluorescence intensity of the NPC-GQDs-PEG after shaking in PBS buffer as a function of time. UV-Vis-NIR spectra were acquired using a PerkinElmer Lambda 750 UV/Vis spectrophotometer. Magnetic properties were investigated using a vibrating sample magnetometer (Lake Shore 7307). *T*2-weighted MRI relaxation and image acquisition were with a 3 T clinical MRI scanner (Philips). The PAI experiments were performed using a multispectral photoacoustic tomography system (MSOT inVision).

### Cell cytotoxicity assay and *in vitro* confocal imaging

L929 murine fibroblast cells (ATCC) and MCF-7 human breast cancer cells (ATCC) were used for *in vitro* experiments. Cells were cultured in ATCC recommended standard conditions. MCF-7 and L929 cells were seeded in 96-well plates and incubated first for 24 h at 37 °C under 5% CO2. After the cells were incubated with various concentrations of NPC-GQDs-PEG for 24 h, 48 h, and 72 h, the standard cell counting kit (CCK8) (Sigma-Aldrich) assay was performed to test the cell viability. The MCF-7 cells were co-cultured with NPC-PEG (100 μg/ml) and NPC-GQDs-PEG (100 μg/ml) for 4 h and then imaged by the confocal laser scanning microscopy (CLSM, Zeiss LSC710META) to study the intracellular targeting effect. Cell nuclei were stained with 4’,6-diamidino-2-phenylindole (DAPI), and cytoskeletons were stained with Rhodamine B (red)-labeled phalloidin (Sigma).

### Animals

All *in vivo* experiments were approved by the Animal Care and Use Committee (IACUC) of the Chinese Academy of Medical Sciences Tumor Hospital (#NCC2019A010). Strict adherence to animal care and use protocols was ensured throughout the study. BABL/c nude mice (about 5 weeks old) were purchased from Beijing Vital River Laboratory Animal Technology Co., China. Mice were implanted MCF-7 cells (2 × 10^6^, 0.2 ml in H-DMEM culture medium without FBS) subcutaneously near the right side of the axillary. The mice were used when their tumor volumes approached about 60-80 mm^3^.

### Blood clearance and *in vivo* measurement of NPC-GQDs-PEG and GQDs in tumor and organs

12 Female BALB/c mice were randomly grouped (n=4): GQDs(-OH), GQDs(-NH2), and NPC-GQDs-PEG. Each formulation (0.2 ml per injection) was injected intravenously at 20 mg kg^−1^ for the NPC-GQDs-PEG group 5 mg kg^−1^ for GQDs groups. At timed intervals, 10 μl blood samples were collected from the tail veins mice at certain time intervals post-injection of NPs and dissolved in 1 ml of the lysis buffer (1% SDS, 1% Triton X-100, 40 mM tris acetate). The concentration of NPC-GQDs-PEG in blood was determined with inductively coupled plasma mass spectrometry (ICP-MS) by analyzing Fe concentrations after the samples were digested with digesting solution (HCl: HNO_3_ of 3:1 by volume). For GQDs(-OH) and GQDs(-NH2) groups, 10 μl blood was drawn and dissolved in the 1 mL lysis buffer and determined from the fluorescence spectrum (F-7000, Hitachi, Ex/Em: 490nm/550 nm) by comparing with a standard curve. The standard curve was obtained by measuring the fluorescence spectrums of series GQDs solutions with known concentrations. Additionally, a blank blood sample without GQDs was measured to determine the blood auto-fluorescence level, subtracted from the fluorescence intensities of injected GQDs during the concentration calculation. The decay curve in the blood was fitted with a two-compartmental model to extract blood half-time using a Microsoft add-in tool. For real-time quantification of NPC-GQDs-PEG or GQDs in the tumor, four tumor-bearing mice were intravenously injected with 0.2 ml NPC-GQDs-PEG solutions (20 mg kg^−1^) or GQDs-saline solution (5 mg kg^−1^). Then, a small tissue sample was resected from the tumor before and after intravenous injection at different time points. After quantitative weighing, these small tumor tissues of the NPC-GQDs-PEG group were dissolved in the digesting solution (HCl: HNO_3_ of 3:1 by volume). The concentration of the NPC-GQDs-PEG was determined with ICP-MS by analyzing Fe concentrations. For GQDs groups, a small sample tissue was resected from the tumor, weighed, and homogenized in the lysis buffer (the same as the above). Clear homogeneous tissue solutions were diluted 100 times to avoid light scattering and self-quenching during fluorescence measurement. The measurements were repeated five times following the above methods to ensure reproducibility and accuracy. Compared with the standard curve of GQDs, the concentration of GQDs in tumor tissue was known.

In the quantitative biodistribution analysis of major organs, the major organ tissues were collected from the mice, weighed, and solubilized by a lysis buffer at different time intervals post-injection. The homogenized tissue lysates were diluted and measured by fluorometry (GQDs) or ICP-MS (NPC-GQDs-PEG) to quantitatively determine the concentrations of NPs.

### Real-time monitoring of the dynamics of NPC-GQDs-PEG in tumor and major organs local by Fibered confocal fluorescence microscopy

The real-time monitoring of NPC-GQDs-PEG *in vivo* was performed using a fibered confocal fluorescence microscopy (FCFM) imaging system (Cellvizio®, Mauna Kea Technologies). The device was equipped with two flexible fiber microprobes. One is the S-1500 probe (0.3 mm of tip diameter) with a penetration depth of 15 μm below the tissue surface. The other is the Ultraminio microprobe with high resolution (2.6 mm diameter, 1.4 μm spatial resolution). An animal operation apparatus, mainly a stereotaxic instrument and reversed binocular microscopy, was introduced to fix and locate the mice during operation. MCF-7 tumor-bearing BALB/c mice were injected via a tail vein with 150 μl NPC-GQDs-PEG (25 mg kg^−1^). Then, 100 μl Evans Blue (2% wt.) was injected intravenously as a microvascular contrast medium at least 4 hours before FCFM imaging. For local tumor imaging, a small incision (4 mm) of tumor epidermis was made to provide access for imaging the surface and deep tumor tissue by the S-1500 probe (S-probe) and Ultraminio microprobe (D-probe), respectively. The exposed tissues were kept moist during the acquisition time with PBS buffer pre-warmed to 37 °C. FCFM imaging was performed before, 2, 12, 24,36, and 72 h post-injection. The dual-band excitation/emission allowed the simultaneous recording of the fluorescence signal from NPC-GQDs-PEG (488 nm, green) and Evans blue (660 nm, red). Furthermore, the quantitative fluorescence intensity of the NPC-GQDs-PEG to Evans blue signals was quantified using Image J software.

Similarly, the FCFM imaging of some major metabolic organs was performed like the FCFM imaging of tumors. To better conduct the imaging, mice were placed in the supine position during acquisition. After anesthesia, a small incision (about 4 mm) was made to provide access to put the flexible confocal microprobe (D-probe) into the abdominal cavity. FCFM imaging was performed before, 8, 24,36, and 72 h post-injection. After imaging each time, the belly was sutured. Additionally, 10 ml/kg body weight buprenorphine solution was s.c. They injected once postoperatively before the animal woke up. They were repeating, if necessary, at any point during daily postoperative in line with all relevant guidelines and regulations of IACUC. The quantitative fluorescence intensity of the NPC-GQDs-PEG to Evans blue signals was quantified using Image J software.

### MRI/PAI multimodality imaging *in vivo*

For MRI and PAI, BALB/c mice bearing MCF-7 tumor were intravenously injected with 150 μl NPC-GQDs-PEG (25 mg kg^−1^) and then observed by using a clinical 3T MRI scanner (Philips) or a multispectral photoacoustic tomography system (MSOT inVision) at various time points, including before treatment and post-12 h, 24 h, 48 h, and 96 h.

### Fluorescence imaging-guided surgery and histological analysis

Fluorescence imaging-guided (FMI) surgery for breast cancer tumors was performed using our self-built navigation system, and the core device is a stereo fluorescence microscope (LeicaM165FC/205FA). A total of 18 mice were randomly divided into three groups: (1) control group: no treatment; (2) N-surgery group: normal surgery; (3) FMI-surgery group: fluorescence imaging-guided surgery after intravenous injection with NPC-GQDs-PEG (25 mg kg^−1^). Surgeons were asked to remove tumor tissue with maximum normal tissue protection based on visual inspection and palpation. Color and fluorescence images were simultaneously acquired and dynamically displayed. In addition, fluorescence images were acquired during the procedure to determine whether small residual tumor tissues still existed. If microtumors remained, the surgeons continued the resection under the fluorescence imaging until all residual microtumors were removed. Three experienced surgeons were asked to conduct the surgery simultaneously. After the surgery, the resected tissue was histologically examined, and the survival condition of mice in different groups were continuously monitored for 9 weeks.

### Statistical analysis

The data are presented as means ± standard deviation, as noted in each case. The number of samples, n, is provided. A two-way analysis of variance (ANOVA) was used.

## Data availability

The data that support the figures within this paper is available upon reasonable request.

## Acknowledgments

The authors thank the National Natural Science Foundation of Youth Program of China (No. 81901813) and Key Social Development Project of Hainan Provincial Department of Science and Technology of China (ZDYF2020137) for support of funding.

## Author contributions

H.Y., X-D.S., J.T., and F-Y.K. discussed, conceived the project. H.Y., J-Y.W., and Z-Y.X. fabricated and characterized the NPC-GQDs-PEG nanocomposite. H.Y., Q.W., W-T.S., and L-Y.Z. designed the targeting function, performed *in vitro* experiments. H.Y. and L-Y.Z. established the mouse model. H.Y., and. Q.W. performed *in vivo* blood clearance and tumor accumulation experiments. H.Y. performed *in vivo* experiments of real-time tracking NPC-GQDs-PEG in local tumor and organs. H.Y., and W-T.S. conducted fluorescence imaging-guided surgery and *in vivo* multimodality imaging. H.Y. prepared figures and wrote the manuscript, S-S.F. and F.Y.K. revised it.

